# Fusion gene discovery in single cells from high throughput long read single cell transcriptomes

**DOI:** 10.64898/2026.01.13.699333

**Authors:** Cheng-Kai Shiau, Hsiao-Yun Lin, Timothy Pan, Xiaodong Lu, Minhua Wang, Yueying He, Lina Lu, Jindan Yu, Ruli Gao

**Author notes:** Correspondence (R.G.).

## Abstract

High throughput long-read single cell RNA sequencing allows for fusion gene discovery in thousands of cells in parallel, which is however constrained by the lack of robust computational tools. Here, we develop LongFUSE that employs the XOR logic operation and stringent filtering criteria for the accurate detection of cell-specific fusion genes and their splicing isoforms. LongFUSE outperforms existing tools in both simulation and real data, demonstrating high accuracy and robustness.

## Main

Gene fusion plays critical roles in driving tumorigenesis^1-4^. Historically, fusion gene analysis has been limited to bulk levels, resulting in a mixed understanding of their oncogenic functions and activation models due to cellular heterogeneity. The emerging long-read single cell RNA sequencing (LR-scRNAseq) enables direct sequencing of full-length fusion transcripts in thousands of cells in parallel. Existing computational tools for fusion gene detection^5-8^ showed high false discovery rates due to inadequate consideration of the analytical challenges in LR-scRNAseq, such as the existence of chimeric reads derived from mis-ligated fragments (*e*.*g*., chimera artifacts)^9, 10^, transcriptional read-through^11^, *trans*-splicing^12^ and the large amount of cell-to-cell variations.

To address these limitations, we develop LongFUSE, a robust computational tool for the accurate detection of fusion genes and their splicing isoforms in individual cells from LR-scRNAseq data. LongFUSE devises XOR logic circuit to achieve fast identification of fusion candidates among large numbers of high-softclipping reads, followed by a selection process for true fusion genes with stringent filtering criteria. Critically, LongFUSE rectifies chimera artifacts, mitigating false discoveries of fusion events and erroneous assignment of cell identities. Moreover, LongFUSE detects exact genomic breakpoints by examining reads of nascent RNAs and assembles potential splicing isoforms of fusion genes from reads mapped to mature RNAs. Lastly, LongFUSE quantifies the expression levels of fusion genes and their splicing isoforms in individual cells with unique molecular identifiers (UMIs). These endeavors ensure the precise discovery of fusion genes at both single cell and single molecule levels.

The computational workflow of LongFUSE starts from scanning and curation of false-ligated chimeras that often occur during the library preparation steps of long read sequencing^9, 10^ (**Fig 1a**). After curation, all singleton reads harboring true cell barcodes and unique molecule identifiers are re-aligned onto the reference genome (GRCh38) using minimap2^13^, followed by XOR logic operation to select candidate reads that harbor fusion genes (**Fig. 1b-c**). In this process, the multiple mapping results against reference genome of each high-softclipping read were aggregated and summarized into a unified binary string using XOR logic circuit (**Online Methods**), which simplifies conditional logic looping and favors speedy searching among large numbers of long-read data. To select candidates for fusion-harboring reads, we derived two parameters, XOR coverage and XOR gap, based on the unified binary string. Reads with high XOR coverage and low XOR gap are expected to be derived from fusion transcripts (**Fig 1b**), whereas reads with low XOR coverage and/or high XOR gap are likely due to low genomic complexity (*e*.*g*., repetitive insertions of transposable elements into multiple genomic regions) (**Fig 1c**). Next, we compiled a collection of filters (**Online Methods**) to remove false discoveries from non-quantitative factors, such as chimeric readouts of read-through events or fusion partners which were homologous. Lastly, the list of confident fusion gene pairs in each cell are characterized through quantification of expression levels, detection of genomic breakpoints and assembly of splicing isoforms (**Online Methods**) to support a range of downstream isoform analyses across cell populations.

**Figure 1.**
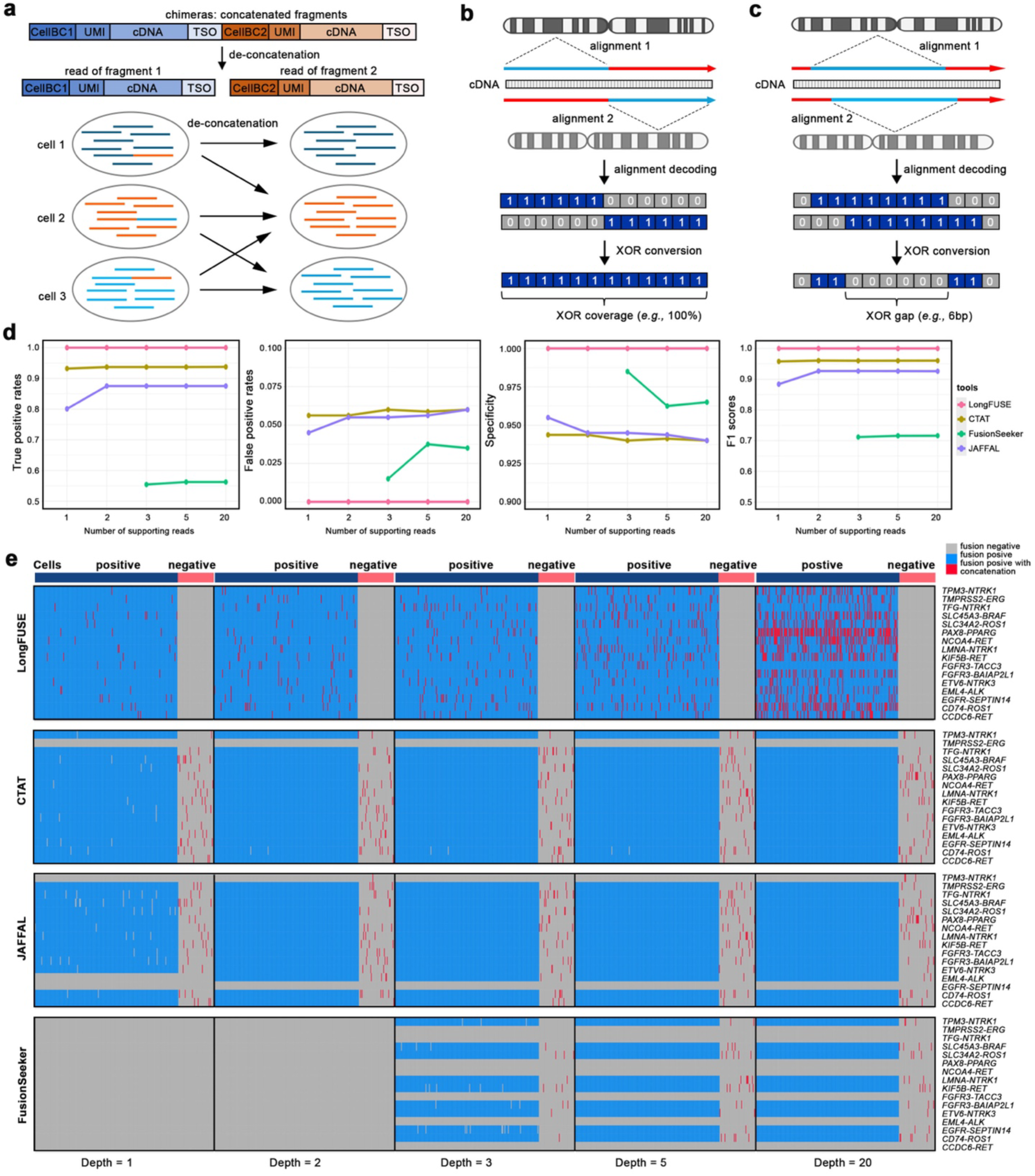
Development of LongFUSE workflow and benchmarking its performance. **a**, Curation of chimera artifacts. XOR selection for candidate fusion genes (**b**), and repetitive low complexity elements (**c**). **d**, Comparison of LongFUSE performance with three existing tools. **e**, Heatmap of 16 spike-in fusion genes in single cells across depths and three existing fusion detection tools.

To benchmark the performance of LongFUSE, we simulated fusion-harboring LR-scRNAseq data by spiking in 16 chemically synthesized fusion transcripts^8^ into our in-house LR-scRNAseq data of a human PBMC sample at different depths (**Online Methods**). In addition to fusion-harboring reads, we spiked in a portion of synthetic chimera reads into a subset of cells to investigate false discovery due to sequnecing artifacts. Next, we applied LongFUSE and three leading fusion detection tools, including CTAT-LR-fusion^8^, JAFFAL^5^, and FusionSeeker^14^ onto these data to perform fusion gene discovery in individual cells. Comparative analyses revealed that LongFUSE outperformed all these tools (**Fig. 1d**) and successfully detected all spike-in fusion genes in all cells across various sequencing depths and rescued the true cell identifiers from the chimera reads (**Fig. 1e**). All three existing tools showed a low capability in rescuing chimea artifacts, leading to false discovery of fusion events and erroneous assignment of cell identifiers. Both CTAT-LR-fusion^8^ and JAFFAL^5^ showed reasonable performance, with CTAT-LR-fusion^8^ failing to detect *TMPRSS2-ERG* and JAFFAL^5^ failing to detect *TPM3-NTRK1* and *EGFR-SEPTIN14* (**Fig. 1e**). As a side note, CTAT-LR-fusion^8^ had initially detected *TMPRSS2-ERG* harboring reads, but was filtered out by the default criteria, resulting in the false negative discovery of a critical fusion event. Together, these data highlight the robustness and accuracy of LongFUSE in detecting true fusion genes in single cells across various sequencing quality conditions.

Next, we applied LongFUSE onto an in-house LR-scRNAseq data of a prostate cancer cell line, VCAP, known for harboring the *TMPRSS2-ERG* fusion. The sequencing data was generated from fresh cell cultures using scNanoRNAseq, as previously published^15^. On average, we obtained 26,983 consensus reads per cells, among which 3,775 showing high-softclipping alignments were subjected to LongFUSE workflow. Remarkably, chimera reads concatenating two or three random fragments accounted for large portions (63% ± 5% SD) of high-softclipping reads (**Table S1**), while the percentages over total raw reads appeared lower (9% ± 0.5% SD) (**Fig. 2a**). These data emphasized the critical importance of distinguishing chimera artifacts in long-read single cell data analysis. After curation, all singleton reads were re-mapped onto the reference and subjected to XOR operation to identify fusion candidates. The binned XOR coverage showed a punctuated drop around 90%, suggesting a potential cutoff for fusion positive events. Consistently, the XOR gap dropped to 10 nucleotides when XOR coverage increased above 90% (**Fig. 2b**). To balance the tradeoff between sensitivity and specificity, we set the XOR coverage parameter to be greater than 80% and XOR gap to be less than 30 nucleotides as the initial criteria for selecting fusion candidates. Noteworthy, this step yielded a 60-fold enrichment for fusion harboring reads from the high-softclipping read pools (**Fig. 2c-d, Table S1**), highlighting the robustness of the XOR-based strategy in selecting true fusion genes. Next, fusion candidates were subjected to a series of filtering criteria to remove additional false discoveries related to non-quantitative factors, such as read-through events and homologous gene annotation artifacts. Collectively, the miscellaneous collection of filtering steps demonstrated a 6-fold enrichment for true fusion gene pairs in these cells (**Fig. 2c-d, Table S1**).

**Figure 2.**
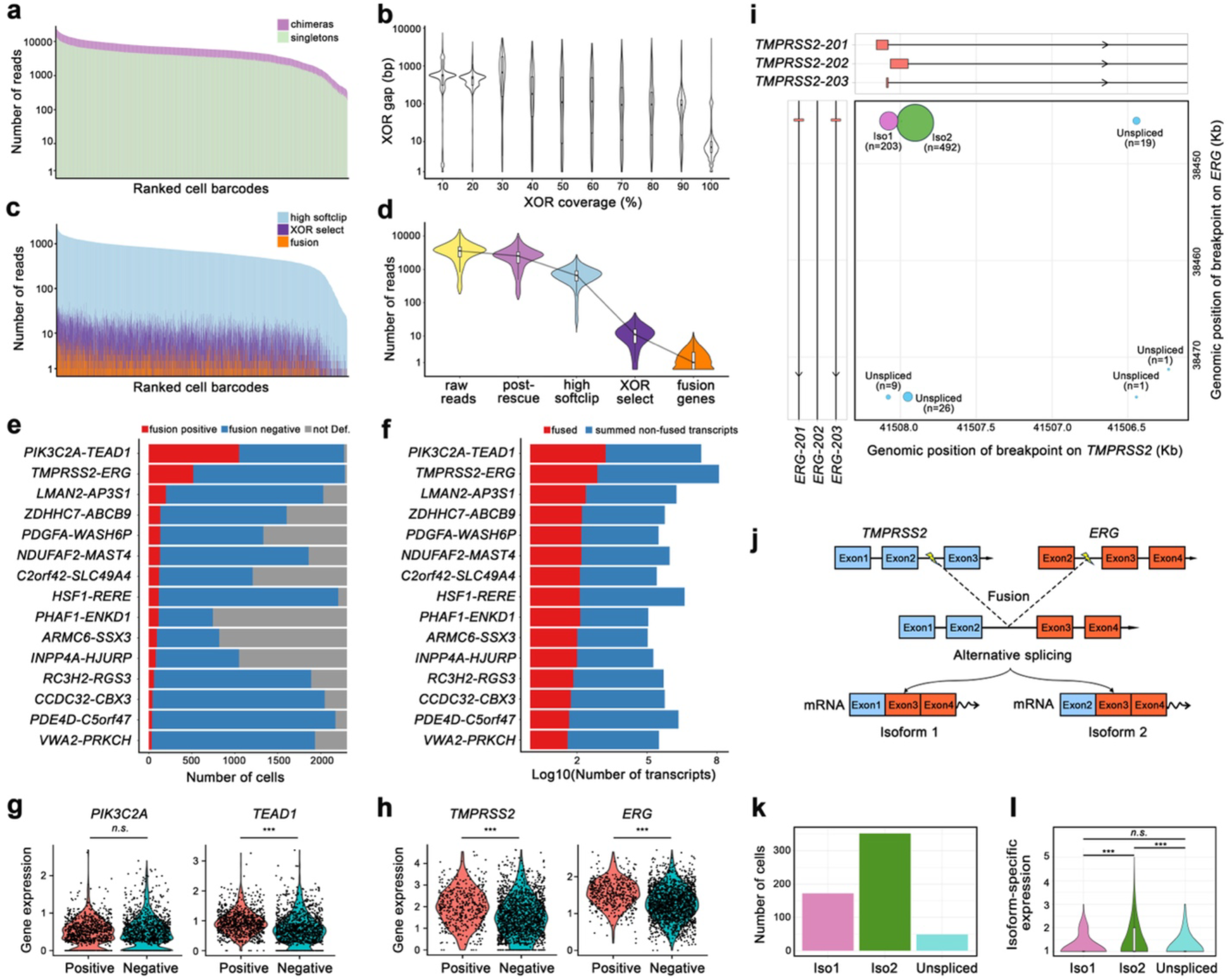
Single cell fusion gene discovery in VCAP cell line. **a**, Composition of raw singleton and chimera reads in single cell. **b**, Distribution of XOR coverage and gap. **c**, Composition of post-curation read types. **d**, Sequential enrichment for fusion-harboring reads. **e**, Numbers of cells expressing fusion genes. **f**, Number of fusion transcripts and the summed wild-type transcripts. **g-h**, Expression levels of top two pairs of fusion partners. **i**, Supporting reads aligned to exact genomic positions. **j**, Inference of splicing isoforms. **k**, Cellular prevalence and **l**, expression levels of isoform-specific fusion transcripts.

To facilitate analysis at cellular levels, LongFUSE classified fusion-positive cells based on the existence of fusion-harboring reads and differentiated fusion-negative cells from ‘not defined’ cells by cross-checking the presence of wild-type transcripts of either fusion partners. Data showed that the *PIK3C2A-TEAD1* fusion was detected in 1,050 cells (*i*.*e*., 38.5% cellular frequency), ranking it as the top expanded fusion event in this cell line, followed by the *TMPRSS2-ERG* fusion that was detected in 516 cells (*i*.*e*., 18.9% cellular frequency) (**Fig. 2e**). Of note, the expression levels of fusion transcripts in fusion-positive cells were lower than the summed expression of both fusion partners in fusion-negative cells (**Fig. 2f**), but higher than the expression of individual fusion partners (**Fig. 2g-h**) with statistical significance (Wilcox *P-value*<0.001) except for *PIK3C2A* (Wilcox *P-value* =0.09). These results showed-cased the unique power of LongFUSE in studying the exact oncogenic dosage effects of fusion genes in single cells.

Next, we performed a detailed characterization of the *TMPRSS2-ERG* fusion that was broadly reported in prostate cancer^3, 16, 17^. As expected, the precise breakpoints were detected in the intronic regions between exons 2/3 of *TMPRS2* and exons 2/3 of *ERG* genes, supported by a total of 56 pre-spliced reads (**Fig. 2i**). Interestingly, most reads were mapped to splice junction regions with less than a 5-nucleotide distance to the closest exon boundaries, allowing for isoform inference of the fusion transcripts. In total, 203 reads were mapped across the junctions between exon 1 of *TMPRSS2* and exon 2 of *ERG*, while 492 reads covered junctions between exon 2 of *TMPRSS2* and exon 2 of *ERG* (**Fig. 2i**). These reads were assembled into two major splicing isoforms (Iso1 and Iso2) that resulted in distinct coding sequences of the *TMPRSS2-ERG* fusion gene (**Fig. 2j**). The two isoforms showed differential cellular prevalence, with 203 cells having utilized Iso1 contrasting to 352 cells that utilized Iso2 (**Fig. 2k**). Furthermore, Iso2 demonstrated significantly higher expression compared to Iso1 (Wilcoxon *P-value*=6.13e-4). Together, these results exemplify the unique strength of LongFUSE in supporting the detailed analysis of fusion genes at both cellular and isoform levels.

In summary, we report a new and robust computational tool called LongFUSE for the detection of fusion genes and their splicing isoforms in individual cells from high throughput LR-scRNAseq data. LongFUSE outperforms existing tools, achieving high accuracy and robustness through XOR logic operations and stringent filtering criteria. Importantly, it precisely reports coding gene pairs, genomic breakpoints, expression levels, and splicing isoforms of fusion transcripts in single cells, facilitating comprehensive downstream analyses of fusion genes across heterogeneous cell populations.

## Data Availability

The raw and processed scNanoRNAseq data generated in this study are deposited and freely accessible in the Gene Expression Omnibus (GEO) database under accession number GSE310978. The simulated fusion containing LR-scRNAseq data are deposited in Zenodo and can be accessed at: https://zenodo.org/records/17677124. The bulk long-read data of synthetic fusion transcripts were downloaded from Sequence Read Archive (SRA) under PRJNA1076207 (SRR27957791, SRR27957792, and SRR27957793).

## Code Availability

All code and detailed documentation of LongFUSE methods are released at GitHub (https://github.com/gaolabtools/LongFUSE).

## Acknowledgments

We thank research funding supports provided to R.G. by the National Institute of General Medical Sciences (NIH R35GM142539) and the National Heart, Lung, and Blood Institute (NIH 1R01HL160552).

## Author Contributions

C.K.S. led the development of LongFUSE, performed the data analysis and wrote the manuscript. H.Y.L and M.W. performed sequencing experiments. T.P., X.L., Y.H., L.L., and J.Y. reviewed data, participated in discussions, and edited the manuscript. R.G. supervised the overall study, conceived the concepts, designed the computational tool, analyzed the data, and wrote the manuscript.

## Online Methods

### Cell line and tissue samples

The VCaP cell line was purchased from ATCC (cat. no. CRL-2876^™^). The PBMC sample was collected from a heart transplantation patient at 22-month post-transplantation in Northwestern Memorial Hospital in collaboration with Drs. Jane Wilcox and Arjun Shina. All activities of this study complied with relevant ethical regulations of Institutional Biomedical Review Board at Northwestern University.

### Long-read single cell RNAseq and data processing

The long-read single cell RNAseq was performed using scNanoRNAseq method as we recently published^15^. To do so, frozen cell stocks were thawed and resuspended in cell culture medium. After cell filtering and counting, the single cell suspensions were loaded onto single cell capture and barcoding platform (10X Genomics, Chromium iX, Single Cell 3’ v3.1 protocol) to attach cell barcodes (CellBCs) and unique molecular identifiers (UMIs) onto poly(A) tailed mRNAs of single cells. Barcoded cDNAs were then enriched for library preparation using LSK114 kit (Oxford Nanopore Technologies, ONT), and then sequenced on our in-house long read sequencer (PromethION CapEX24, ONT) using R10.4 Q20+ flow cells, with one sample per flow cell. After sequencing, the raw sequencing signals were base-called using Dorado (ONT) in super-accurate mode) and stored in FASTQ format. The raw master FASTQ files were subjected to scNanoGPS for demultiplexing into single cells and single molecules and mapping onto reference genome. The reads harboring fusion genes are contained in high-softclipping reads that showed high mappability onto multiple genomic regions. Single cell high-softclipping reads were extracted from the mapped BAM files and subjected to LongFUSE workflow.

### De-concatenation of chimera reads of ligated fragments

Long-read sequencing often generated unneglectable chimera reads of concatenated DNA fragments that were caused by mis-ligation during library preparation steps^9, 10^. The chimera reads contain multiple sets of barcodes, show high-softclipping mapping results and appear as harboring fusion genes, causing significant amounts of false discovery in fusion gene detection. Leveraging the expected patterns of barcode and adaptor sequences, LongFUSE identified and broken down the chimeras into singleton reads, as such each full-length reads only contain one cDNA fragment along with the true cell barcode and unique molecular identifiers. After curation, all singleton reads were re-aligned against the reference genome with minimap2^13^. The singletons with high-softclipping mapping results were then extracted and used for downstream detection of fusion transcripts in single cells.

### Selection of fusion candidates with XOR logic operation

To improve the speed and simplify logic looping, the XOR logic circuit principles were adopted to select candidate fusion-harboring reads among high-softclipping reads. To do so, the two mapping-quality strings corresponding to the top two mapped regions were first converted to individual binary strings, where highly mappable loci were given 1 or 0 otherwise. Next, the two binary strings of each read were unified to generate a new string through exclusive OR (XOR) logic gate computing, in which 1 was assigned to a position if the values on the two initial strings were different or 0 if the two initial values were same (both 0, or both 1). As such, the multiple alignment conditions and outcomes were aggregated and summarized into a unified binary string, which 1 represented high mappability of only one genomic region and 0 represented several known and unknown possibilities, for instance, both high or both low mappability onto the top two genomic regions or the certain loci had low data quality. This XOR operation simplified conditional looping and greatly favored speedy search for fusion candidates among large numbers of long-read data. Next, we derived two parameters to select candidate fusion-harboring reads, e.g., XOR coverage and XOR gap. XOR coverage calculates the percentages of all 1s in the unified logic strings, associating with the confidence of unique mapping to two distinct genomic regions. XOR gap calculates the number of 0s between the boundaries of two consecutive 1s, associating with the continuity between two genomic mapping and potential data quality related insertion and deletions. Based on the empirical observations and trade-off between sensitivity and specificity, XOR coverage ≥ 80% and XOR gap ≤ 30nt were set as the default cutoff and was left open for users to adjust. In our tested data, these settings showed ∼100X enrichment for fusion-harboring reads.

### Miscellaneous filtering steps

To boost speeds, singleton high-softclipping reads from unassembled scaffolds/contigs, mitochondrial chromosomes, and other non-genomic regions were excluded from XOR operation. Reads with discordant alignment directions or high-abundance multiple alignments were also set aside to improve detection confidence. The candidate fusion gene pairs within 3 Kb regions of two adjacent genes were removed to exclude potential read-through events. The candidate fusion gene pairs belonging to paralog/homolog genes according to database were also filtered out to mitigate annotation errors. Lastly, candidate fusion detected in less than certain numbers of cells were filtered out to remove unknown random errors. The default cutoff was set as 3 cells, which was open for users to adjust.

### Simulation of fusion containing LR-scRNAseq data

The fusion containing LR-scRNAseq data were simulated by mixing spike-in fusion reads with an in-house data of a human PBMC sample. The nanopore sequencing data of sixteen chemically synthesized Spike-In fusion transcripts were downloaded from a previously published^8^. The fusion-harboring reads with confident alignment were extracted, attached with barcode sequencing using same library structures as of LR-scRNAseq, and then *in silico* mixed into the PBMC dataset at different read depths (N=1, 2, 3, 5, and 20). Similarly, chimera reads were synthesized to have multiple sets of cDNA fragments and barcodes and then injected into the LR-scRNAseq data at different read depths.

### Benchmarking with existing tools

The performance of LongFUSE was compared with three leading fusion gene detection tools, including CTAT-LR-fusion^8^, JAFFAL^5^, and FusionSeeker^14^ using the simulated datasets. All three existing tools were run under default parameters. The calculation results of the sixteen Spike-In fusion genes in individual cells were compared between the four tools, by calculating true positive rate (TPR), false positive rate (FPR), specificity, and F1 score.

### Detection of exact fusion breakpoints and splicing isoforms

To detect breakpoints, the XOR binary string was computed with AND logic gate separately against two alignment binary strings converted from two fusion partner genes. After AND logic gate computing, the last (right most) 1 was identified as the breakpoint of the upstream (left) alignment (denoted as gene A), while the first (left most) 1 was detected as the breakpoint for the downstream (right) alignment (denoted as gene B). The chromosomal positions of the two breakpoints were computed accordingly. To infer out possible splicing isoform, fusion reads were aligned against provided genome annotation (GTF). All possible isoforms fully matched were reported, along with matching isoform ID and exon ID combinations.

